# Peripheral electrical stimulation augments cerebral collateral circulation if performed within a critical time window

**DOI:** 10.1101/2020.06.08.140582

**Authors:** Ming-Chieh Ding, Aritra Kundu, Colin T. Sullender, Andrew Dunn

**Affiliations:** Department of Neurology, Dell Medical School at The University of Austin at Texas, Austin, TX, USA; Department of Biomedical Engineering, The University of Austin at Texas, Austin, TX, USA

## Abstract

Ischemic stroke is one of the leading causes of death and disability in the world. Recent advances in acute stroke care have dramatically improved clinicians’ abilities to reperfuse occluded blood vessels. With these advances, the importance of adjunctive therapies to supplement or complement reperfusion therapy is receiving greater interest. Cerebral collateral circulation is one of such area that is now gaining greater interest in acute stroke care. In this study, we investigate the use of peripheral electrical stimulation to induce functional hyperemia in a mouse animal model in the setting of acute stroke. Using a laser speckle contrast imaging system, we evaluated the use of peripheral electrical stimulation at 1 hour and 3 hours after stroke induction. Results demonstrated that stimulation initiated 1 hour following stroke significantly increase collateral cerebral blood flow, while stimulation at 3 hours after stroke had no appreciable effect. These results suggest that augmentation cerebral collateral circulation may be possible in the setting of acute stroke although there may be a critical time window in which this would have to be initiated.

## Introduction

Ischemic stroke is a neurological injury that results from disruption of blood flow to the brain and is one of the leading causes of death and disability in the world ^1^. Traditionally, acute therapeutic intervention has largely been limited to intravenous thrombolysis with the fibrinolytic medication, recombinant tissue plasminogen activator (tPA). Although the efficacy of tPA has been proven in multiple clinical trials, with tPA being a strong fibrinolytic agent, it carries significant risk of causing hemorrhage ^2^. In addition to the numerous contraindications for the use of tPA, there is also a limited window during which it can be administered (no greater than 4.5 hours from the time the patient was last known to be at baseline) ^3^. However, in recent years, the validation of endovascular therapy, with the utilization of perfusion imaging for patient selection, as a viable therapeutic intervention has revolutionized treatment in acute stroke. This has dramatically improved rates of reperfusion and has also substantially extended the time window of intervention to as far out as 24 hours from the time a patient was last known to be well if imaging suggests there is still salvageable brain tissue ^4–8^. One of the major contributors to the severity of injury and duration of penumbra viability is the state of the collateral circulatory vessels ^9–12^. In situations where there are strong collateral vessels, the injury severity may be decreased or the duration of penumbral viability may be enhanced. However, if the collateral circulation is immature or weak, then the ischemia may rapidly progress to irreversible injury ^13^. Even though the importance of collateral circulation has long been recognized, it remains relatively underexplored as a potential therapeutic target and the general sentiment has largely been that some people are fortunate to have strong collateral vessels and others are not. However, there may be ways to temporarily recruit collateral vessels. Although temporary recruitment may have previously had limited use when the primary vessel frequently remains occluded, now that endovascular therapy has led to higher rates of acute revascularization, even temporary methods of enhancing collateral circulation may have clinical utility and such techniques should be investigated.

One potential method is to utilize functional hyperemia or the concept that cerebral blood flow is closely tied to neuronal metabolic activity ^14^. One method in which functional hyperemia can be modulated in a targeted manner is via peripheral stimulation ^15–18^. The use of peripheral stimulation in acute stroke to modulate injury severity has been explored to a degree in prior work ^19–21^. However, previous studies have not necessarily focused on cerebral blood flow and used surrogate measures of function (e.g. evoked potentials) or used interventions that were not clinically feasible (e.g. initiation of stimulation within minutes of stroke onset) or were in rats, which seem to have different tolerance to ischemia compared to mice ^22–24^. This study seeks to specifically demonstrate that cerebral blood flow, namely collateral circulation, can be augmented during the acute period of ischemic stroke which may have potential benefits for translation to clinical therapy as an adjunctive therapy to reperfusion.

## Methods

To investigate the effects of peripheral stimulation on cerebral blood flow, we utilized 12 C57/BL6 male mice (Charles River Laboratories, Wilmington, MA, USA, 22-25 g, n=4 control animals, n=4 stimulation 1-hour post-stroke, n=4 stimulation 3 hours post-stroke). All procedures were approved by the Institutional Animal Care and Use Committee at The University of Texas at Austin and in accordance with National Institutes of Health guidelines. All efforts were made to minimize the number of animals used and their suffering.

### Cranial Window Placement

Cranial window placement followed a very similar protocol as previously described in greater detail by Clark *et al* ^25^. Mice were anesthetized with isoflurane. The level of anesthesia was monitored via respiratory rate and toe pinch response throughout surgery to ensure adequate sedation. A midline incision was made in the scalp and then a portion of skull over the right parietal lobe was removed using a high speed dental drill to expose the somatosensory cortex. Care was taken to leave the dura intact and saline was periodically added to the drill site to clear debris and minimize heating of the skull. Prior to window placement, dexamethasone (2mg/kg, i.m.) was administered to minimize inflammation and post-operative swelling. A cover glass was placed over the skull opening and sealed with cyanoacrylate and dental cement. Animals were allowed to recover for 1 week following cranial window placement prior to any imaging. During the week of recovery, animals were given carprofen daily (2.5 mg/kg, i.p.) both for pain control and to minimize inflammation that can contribute to clouding of the window.

### Imaging and Photothrombosis

Isoflurane has a well-documented effect on cerebral blood flow ^15,26–28^. However, since this study was focused on relative changes in cerebral blood flow with each animal acting as its own control, the absolute effects of isoflurane should be less of an issue. Regardless, a low-dose isoflurane protocol, as described in Munting *et al* was utilized as their study demonstrated that this particular isoflurane protocol appeared to be have the least effect on cerebral blood flow while maintaining cerebrovascular reactivity ^26^. For imaging sessions, mice were anesthetized with isoflurane (2% induction for no greater than 5 minutes with 1.25% maintenance) in oxygen and affixed to a stereotaxic frame. Periodic tail pinch and respiratory rate were also monitored to ensure proper depth of anesthesia. In addition, arterial oxygen saturation and heart rate from pulse oximetry (MouseSTAT Jr.; Kent Scientific, Torrington, CT, USA) were recorded and temperature was maintained at 37 °C with a feedback temperature control system (55-7030, Harvard Apparatus, Holliston, MA, USA). The generation of photothrombosis was almost identical to the method described in Clark *et al* ^25^. In brief, a green diode laser (532 nm, AD-532-100A, AixiZ, Houston, TX, USA) was utilized in conjunction with a digital micro-mirror device to provide patterned illumination (0.14 W/cm^2^) over distal branches of the middle cerebral artery. The targeted branch supplies a portion of the somatosensory cortex and the advantage of this targeted photothrombosis system is that it is able to occlude a vessel with minimal damage to the surrounding parenchyma by limiting the laser light to the target vessel. Rose Bengal (100 µL, 15 mg/mL i.v., Sigma) was administered retro-orbitally and the laser was activated for 15 minutes to generate thrombosis of the targeted vessel. Cerebral blood flow was monitored continuously throughout the experiment using a laser speckle imaging system ^25,29–31^. For laser speckle imaging, red laser light (685 nm, 50 mW, HL6750MG, Thorlabs, Newton, NJ, USA) was relayed to a CMOS camera (acA1920-155um, 1920 x 1200 pixels, Basler AG, Germany) with 2x magnification and acquired at 150 frames-per-second with a 5 millisecond exposure time using custom software as previously described in Sullender *et al* ^32^.

### Peripheral Electrical Stimulation

Animals undergoing stimulation had two subdermal needle electrodes placed in the forelimb contralateral to the region of infarction. Stimulation was delivered via a constant current stimulator (A365, World Precision Instruments, Sarasota, FL, USA). The stimulation protocol was adapted from the protocol utilized in Burnett *et al* ^21^ where peripheral stimulation was examined in a stroke model utilizing rats. Animals were given current stimulation cycles of 5 Hz for 4 seconds followed by 3 seconds of rest before initiation of the next stimulus. Current amplitude was titrated to isolated limb twitch, but typically was between 0.5-1 mA. Control animals did not have stimulation performed. All animals had 5 minutes of pre-stroke baseline recorded. Regions of interest (ROIs) were selected to encompass the visual recording site and each ROI was separated by approximately 300 µm to help establish geographical surface maps of possible core and penumbra. Within the stimulation groups, one set of animals had stroke induced and then had the aforementioned stimulation protocol initiated 1 hour after stroke while the other group of animals had stimulation initiated 3 hours after stroke was induced. Regardless of the cohort, all animals were imaged continuously for 4 hours after stroke induction. Following completion of imaging, animals were allowed to recover for 24 hours and then behavior was evaluated by performing the forelimb extension test. Following behavioral testing, the animals were then euthanized, perfused with paraformaldehyde, and then had their brains harvested.

## Analysis

Regions of interest (ROI) had relative inverse speckle correlation time, which is proportional to surface cerebral blood flow, calculated using custom software ^29^. Following stroke induction, all regions of interest values were normalized to the pre-stroke baseline values to determine relative change in flow for that region following stroke. To evaluate the effects of stimulation, 5-minute periods were averaged for each ROI prior to stimulation and compared to 5-minute period averages at the end of the stimulation protocol. Results for all regions of interest, separated by cohort were plotted on a scatter plot and regression lines were generated for the data points for each group of ROIs. In addition, scatter plots of ROIs for each cohort were plotted individually and compared to the unity line. Each individual animal had regression lines generated for its set of ROIs and the slopes from each of the three groups were compared to each other via unpaired, 2-tailed t-tests.

## Results

As expected, cerebral blood flow decreases after targeted artery occlusion and stabilizes after a variable amount of time. Regions near the occluded vessel tend to show the most significant decrease in cerebral blood flow from their baseline, while surrounding regions show variable amounts of decreased flow, presumably due to variations in their surface or subsurface collateral circulation ^33^. Figure 1 shows a sample image of vessel occlusion from photothrombosis and the resulting stroke. Figure 2 displays ROIs that demonstrate typical temporal trends for the three different cohorts. In Figure 3, all regions of interest in all animals, separated by cohort (control animals, stimulation 1 hour post-stroke, and stimulation 3 hours post-stroke) are plotted to demonstrate how the ratio of their relative cerebral blood flow changes as a result of stimulation. The x-axis shows the ROIs’ pre-stimulation values and they are plotted against their post-stimulation values on the y-axis. Regression lines were generated for each cohort of data points to demonstrate the effects of stimulation on cerebral blood flow. The regression line slope for the control animals was 1.15 (95% CI: 1.02, 1.28). For the animals that received stimulation 3 hours following the initiation of stroke, the regression line slope was 1.05 (95% CI: 0.93, 1.17). The animals that received stimulation starting 1 hour following the induction of stroke had a regression line slope of 2.05 (95% CI: 1.81, 2.29).

**Figure 1:**
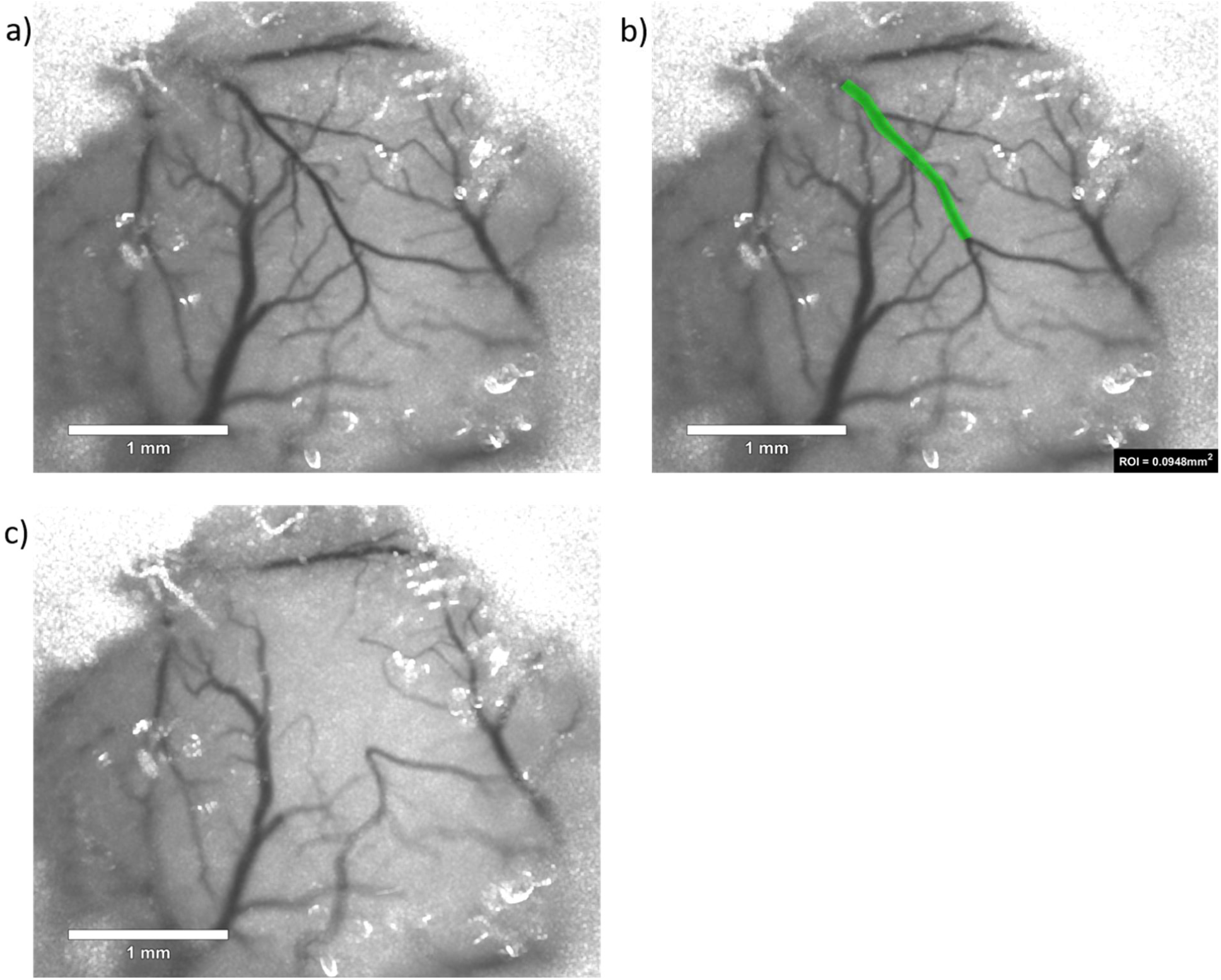
This figure provides a sample image of photothrombosis. a) Displays imaging prior to stroke. b) This image demonstrates vessel targeting with the vessel to be occluded outlined in green. c) This image shows demonstrates occlusion of the target vessel after injection of Rose Bengal and exposure to green laser light with subsequent area of infarct.

**Figure 2:**
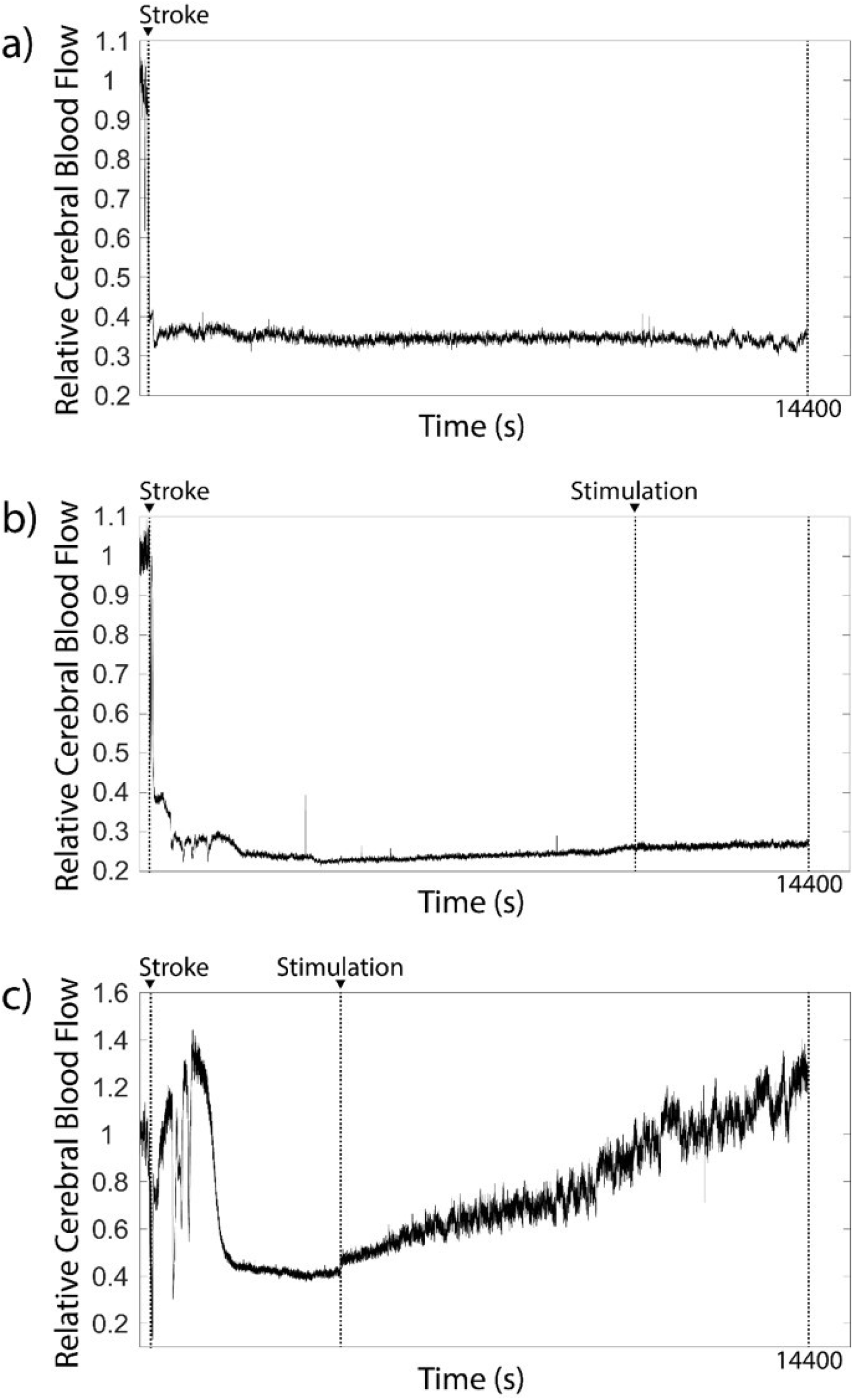
This figure displays an ROI demonstrating a typical temporal progression for each cohort. Sample ROIs were selected from regions approximately 600 µm to 900 µm from the site of vessel occlusion in each animal. The y-axis shows relative cerebral blood flow while the x-axis is the experiment time in seconds. Stroke and stimulation initiation times are labeled. a) The typical progression of cerebral blood flow in a control animal is shown. Following stroke, there is a drop in blood flow that tends to be relatively stable for the duration of the experiment. b) The typical progression of cerebral blood flow in an ROI when stimulation is initiated 3-hours after stroke induction is shown. As expected, there is a decrease in blood flow after stroke induction. When stimulation is initiated, there is no significant change in the cerebral blood flow. c) This figure displays the temporal progression of cerebral blood flow when stimulation is initiated 1 hour after stroke. After stroke induction, there is the expected decrease in cerebral blood flow, but when stimulation is initiated at 1 hour, there is a rapid onset of increase in cerebral blood flow. The time to reach peak blood flow for a given ROI appears to be gradual, but is generally sustainable for the duration of the experiment.

**Figure 3:**
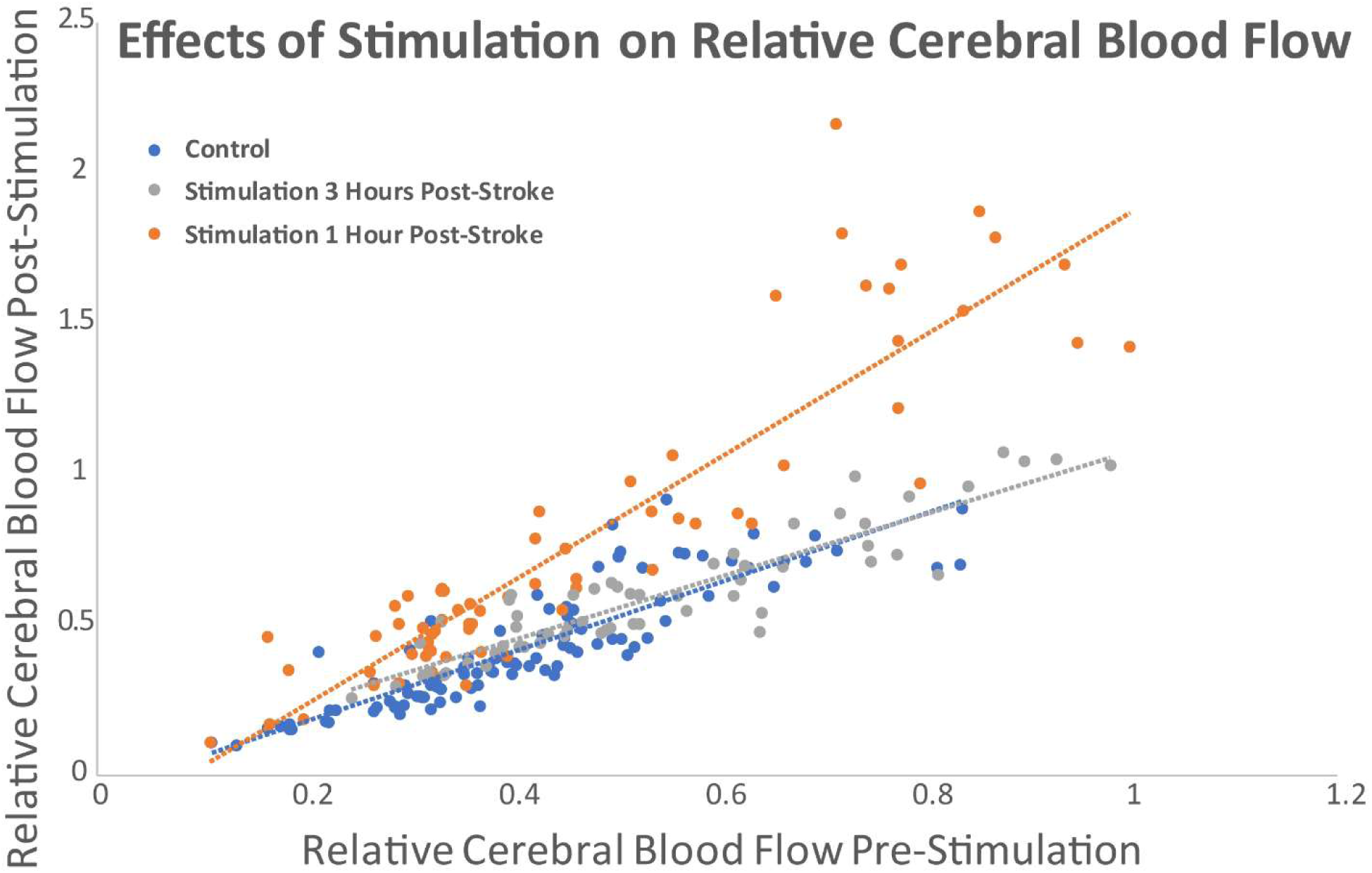
This figure depicts the change in relative cerebral blood for all of the ROIs from all of the animals. The x-axis depicts the relative cerebral blood flow prior to stimulation. The y-axis depicts the relative cerebral blood flow following stimulation. The control (blue) and 3-hour post-stroke stimulation animals (gray) seem to show no significant change in their ROIs as a result of peripheral electrical stimulation. However, the 1-hour post-stroke stimulation animals generally show an increase in their relative cerebral blood flow following stimulation. Regression lines show slopes of 1.15 for the control group, 1.05 for the 3-hour post-stroke stimulation group, and 2.05 in the 1-hour post-stroke stimulation group.

In Figures 4 through 6, the separate cohorts were plotted individually against the unity line with the ROIs from each separate animal delineated and regression lines generated for each separate animal. The average slope of the regression lines for the individual control animals was 0.89 ± 0.14. The average slope of the regression lines for individual animals in the 3-hour post-stroke stimulation group was 0.89 ± 0.22. The average slope of the regression lines for individual animals in the 1-hour post-stroke stimulation group was 1.36 ± 0.20. Unpaired 2-tailed t-tests for the individual animal regression line slopes were performed for the control versus 3-hours post-stroke stimulation group (p=0.98), control animals versus 1-hour post-stroke stimulation group (p=0.01), and the 1-hour post-stimulation group versus the 3-hour post-stimulation group (p=0.02).

**Figure 4:**
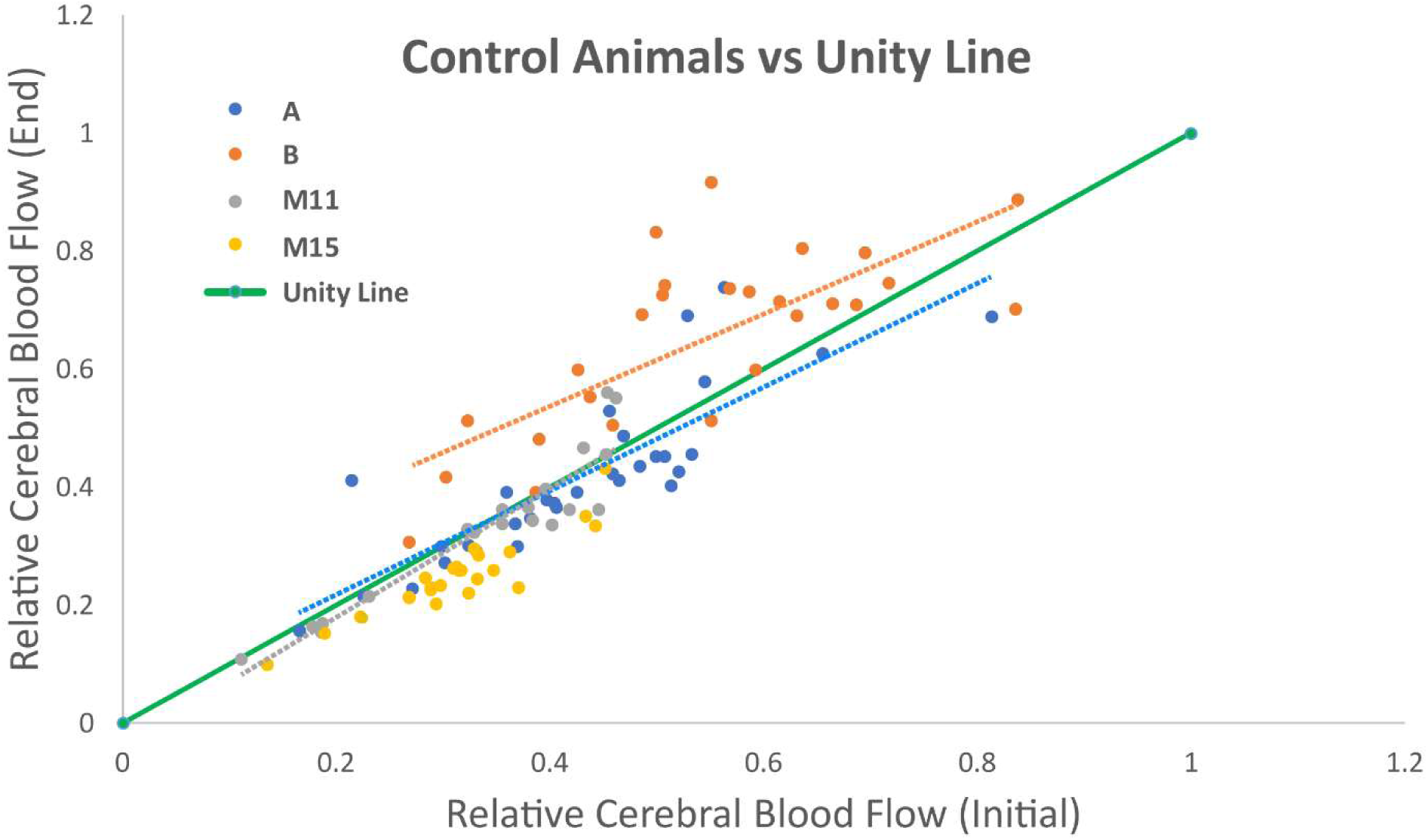
This figure demonstrates the relative cerebral blood flows for the control animal ROIs separated by each individual animal. The x-axis demonstrates the initial relative cerebral blood flow after stroke induction while the y-axis demonstrates the relative cerebral blood flow near the conclusion of the recording. Regression lines for each individual animal are shown. The unity line has been plotted for comparison. The average slope for the regression lines when plotted by each animal individually is 0.89 ± 0.14.

**Figure 5:**
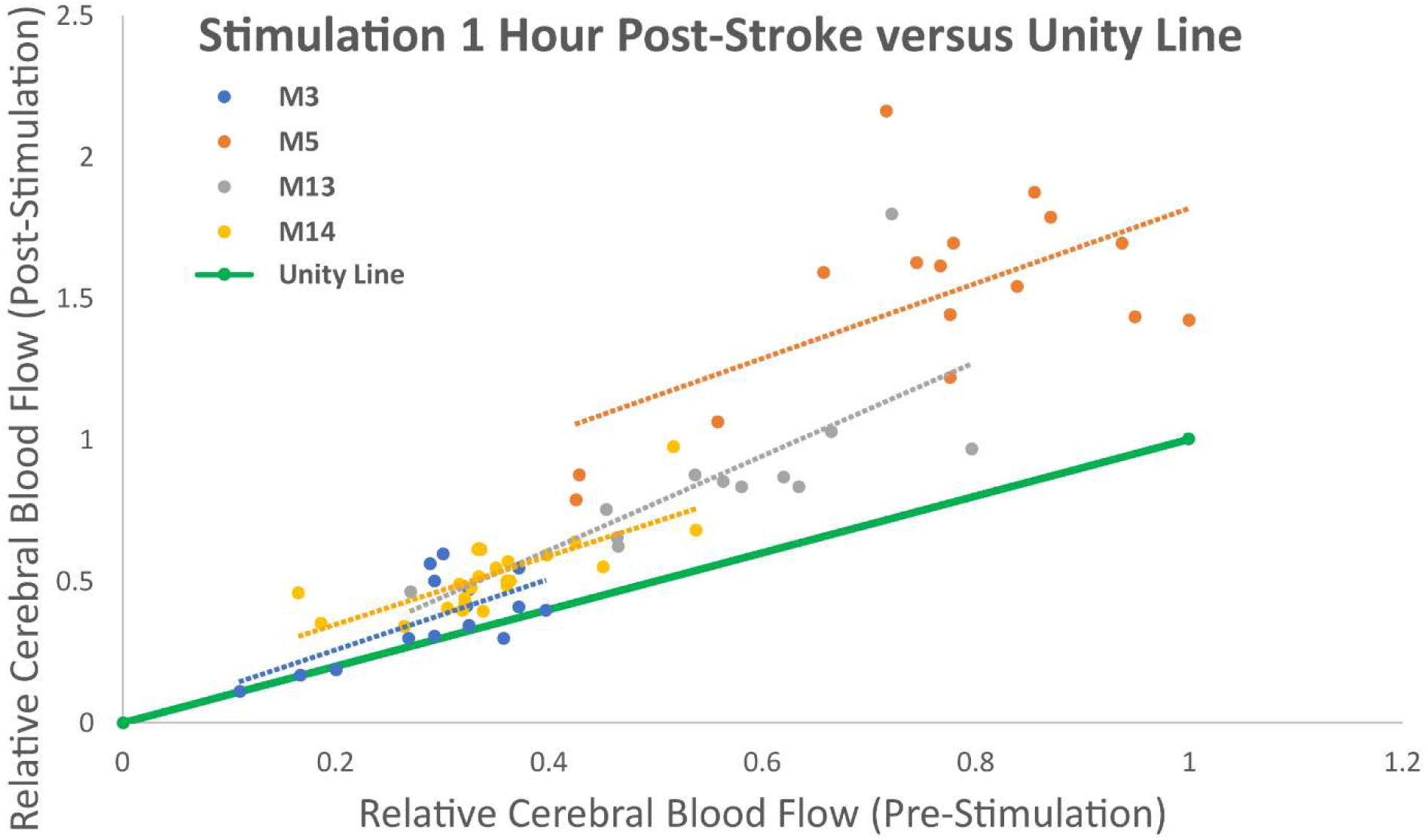
This figure depicts the relative cerebral blood flow for ROIs in the animals receiving stimulation 1-hour post-stroke with ROIs separated by individual animal with the unity line plotted for comparison. The x-axis shows the relative cerebral blood flow in the ROIs just prior to stimulation while the y-axis displays the relative cerebral blood flow in the ROIs at the end of the stimulation protocol. The average slope for the regression lines for the individual animals in this cohort was 1.36 ± 0.20.

**Figure 6:**
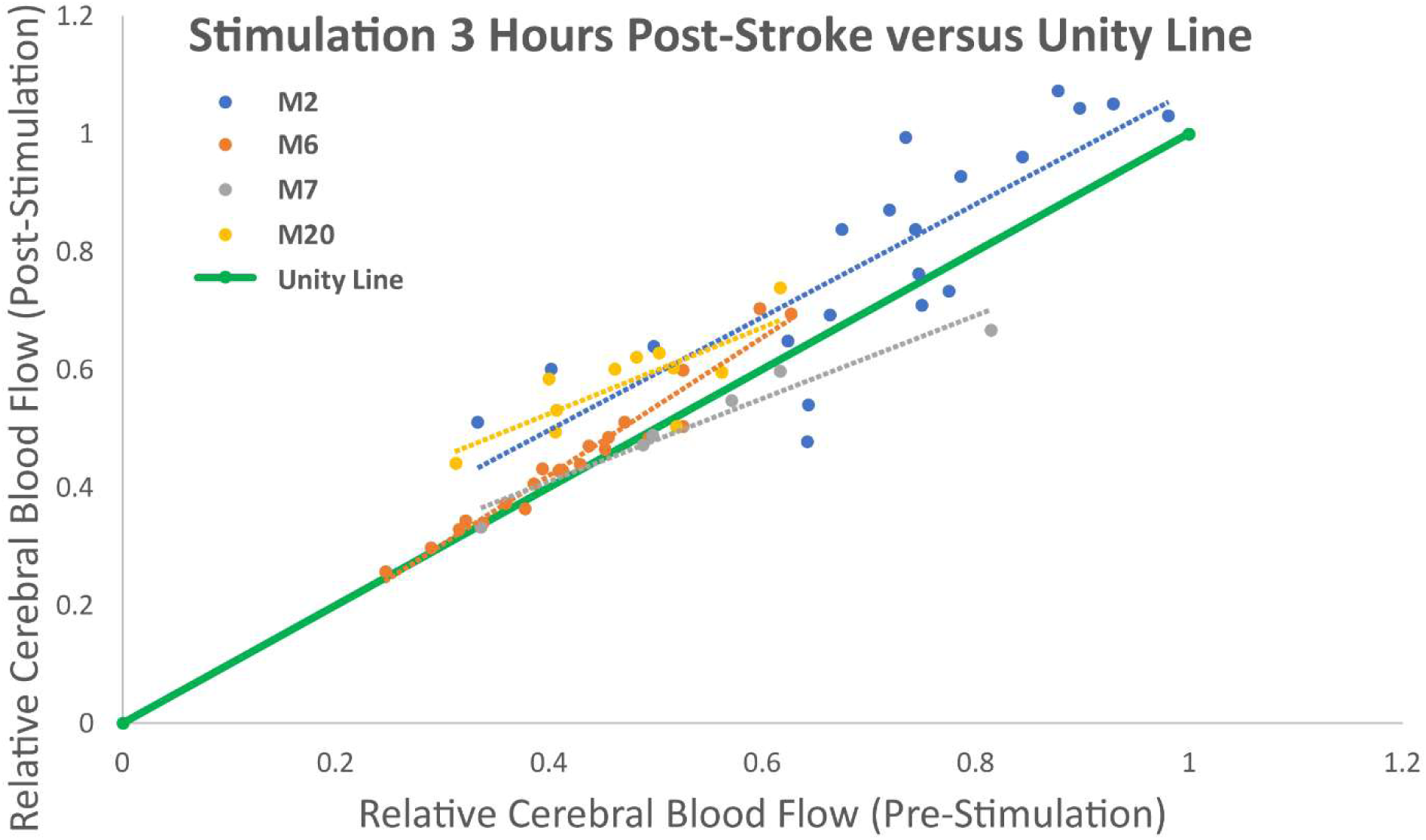
This figure depicts the relative cerebral blood flow for ROIs in the animals receiving stimulation 3-hours post-stroke with ROIs separated by individual animal with the unity line plotted for comparison. The x-axis shows the relative cerebral blood flow in the ROIs just prior to stimulation while the y-axis displays the relative cerebral blood flow in the ROIs at the end of the stimulation protocol. The average slope for the regression lines for the individual animals in this cohort was 0.89 ± 0.22.

24 hours after their stimulation sessions, animals had the forelimb flexion test performed and all animals demonstrated forelimb flexion and rotation towards the side contralateral to the region of stroke when picked up by their tails confirming that even though the injury was limited to a very distal middle cerebral artery branch, it was still sufficient to cause a clinical deficit.

## Discussion

These results appear to demonstrate that peripheral electrical stimulation can potentially modulate cerebral blood flow in the setting of stroke if performed within a specific time window. Our results seem to indicate that when there is no stimulation, cerebral blood flow in regions of interest, as a whole, seem to be fairly stable after a stroke. There is some expected variability and grossly, it seems that regions with post-stroke blood flow above a certain threshold may be more likely to have mild increases in blood flow over time, while below that threshold, regions may have a decline in cerebral blood flow. However, overall, regions tended to be fairly stable, hence the regression line slope of approximately unity. These regional variations are suspected to be secondary to underlying differences in subsurface collateral circulatory vessel distribution. Similarly, the animals that received stimulation 3 hours after the induction of stroke also seemed to show no clear consistent change as a result of stimulation as their regression line slopes, both individually and as a group, did not appear significantly different from the control animal group. These data suggest that stimulation may have no effect on cerebral blood flow after a certain critical time point has passed. Even ROIs that appeared to have a relatively high post-stroke level of cerebral blood flow did not typically demonstrate a significant change in flow as a result of stimulation when initiated 3 hours following stroke. This may suggest by that time point, at least in mice, the injury may be largely static and unable to be modulated from a collateral circulation perspective. It is possible that some tissue may already be irreversibly damaged from a flow perspective or collateral vessels at that point are functioning at maximum capacity or weaker collateral vessels have already collapsed. In the animals that received stimulation one hour after stroke induction, however, the stimulation appeared to consistently result in an increase in cerebral blood flow. Regions that had higher pre-stimulation levels of relative cerebral blood flow generally had more significant increases in relative cerebral blood flow in response to stimulation, but even regions with severe reduction in cerebral blood flow post-stroke demonstrated smaller magnitude increases in cerebral blood flow.

In order to ensure that the results were consistent amongst the cohorts and not the result of a single animal having outlying values for its ROIs, regression line slopes for each animal’s ROIs were also generated. As can be seen from the results shown in Figures 4 through 6, the slopes for the animals in the 1-hour post-stroke stimulation group had statistically significant higher slopes compared to the other two groups, whereas the control animals and the 3-hour post-stroke stimulation group did not have significantly different slopes. Notably, when all ROIs from all animals are analyzed as a single group, the findings are susceptible to being skewed by a single animal outlying animal (e.g. an animal that had spontaneous vessel reperfusion). Hence, the animals were also analyzed individually and this finding was consistent when evaluating all ROIs together and when evaluating the effects of stimulation in each animal individually. Again, this suggests that when stimulation is initiated early, an increase in cerebral blood flow can be generated, but when stimulation is initiated late after injury there is not a clear effect.

Overall, these results suggest that peripheral electrical stimulation may be able to at least temporarily augment cerebral blood flow after stroke if performed within a critical time window. Stimulation performed 1 hour after stroke seemed to consistently cause an increase in cerebral blood flow. These regions could represent areas of functional penumbra as they are still responsive to stimulation and can have flow augmented in response to metabolic demand. It is possible that if flow augmentation can be maintained in these regions until reperfusion, that infarct size, and potentially injury, might be minimized. This effect was not appreciated, however, in animals that received stimulation at 3 hours after stroke induction. The lack of effect of stimulation at 3 hours after stroke induction suggests that there may be a relatively early period after which the injury becomes relatively irreversible or where the collaterals can no longer be augmented.

Our study had several limitations. Since this study utilized a photothrombosis model, which is not reversible in a controlled manner, and had only a transient intervention, histology was not performed as the intervention was not expected to generate a difference in infarct size. However, future studies will be designed with a reversible occlusion model so that histology can be performed and the effects of stimulation on infarct size can be evaluated. Similarly, beyond the forelimb flexion test, in-depth behavioral testing was not performed for similar reasons as a graded behavioral difference was not expected given the transient intervention with a permanent injury. The forelimb flexion test confirmed that the stroke generated a clinically-observable injury, but since the injury model was irreversible, more extensive behavioral testing was not performed. Again, future experiments with a reversible injury model will include more in-depth behavioral testing to evaluate whether transient stimulation can augment cerebral blood flow in a simulated reperfusion model and potentially generate a clinically measurable outcome. Finally, an additional measure of function, such as electrophysiology, will be included in future experiments.

Regardless, these results seem to suggest that collateral circulation can be modulated to some degree, although there may be a certain window during which this may need to be performed. It should be emphasized that although there was a significant difference between stimulation at 1 hour and 3 hours post-stroke in this study, these time points do not necessarily translate to other animals or even other breeds of mice as there is substantial variation in collateral circulation between animals and mouse breeds ^34–37^. Still, this demonstrates that even in an animal that may be potentially more susceptible ischemic insult ^22,38–40^, it appears that collateral circulation can be temporarily augmented in a minimally invasive fashion that could be translated to clinical therapy.

## References

1. Collaborators, T. G. 2016 L. R. of S. Global, Regional, and Country-Specific Lifetime Risks of Stroke, 1990 and 2016. N. Engl. J. Med. 379, 2429–2437 (2018).

2. Group, N. I. of N. D. and S. rt-P. S. S. Recombinant tissue plasminogen activator for acute ischemic stroke. N. Engl. J. Med. 333, 1581–1587 (1995).

3. Powers, W. J. et al. Guidelines for the Early Management of Patients with Acute Ischemic Stroke: A Guideline for Healthcare Professionals from the American Heart Association / American Stroke Association. Stroke vol. 44 (2018).

4. Nogueira, R. G. et al. Thrombectomy 6 to 24 Hours after Stroke with a Mismatch between Deficit and Infarct. N. Engl. J. Med. 378, 11–21 (2017).

5. Marks, M. P. et al. Endovascular Treatment in the DEFUSE 3 Study. Stroke 49, 2000–2003 (2018).

6. Saver, J. L. et al. Stent-retriever thrombectomy after intravenous t-PA vs. t-PA alone in stroke. N. Engl. J. Med. 372, 2285–95 (2015).

7. Campbell, B. C. V. et al. Endovascular Therapy for Ischemic Stroke with Perfusion-Imaging Selection. N. Engl. J. Med. 372, 1009–1018 (2015).

8. Berkhemer, O. A. et al. A Randomized Trial of Intraarterial Treatment for Acute Ischemic Stroke. N. Engl. J. Med. 372, 11–20 (2015).

9. Liebeskind, D. S. Collateral circulation. Stroke 34, 2279–2284 (2003).

10. Nishijima, Y., Akamatsu, Y., Weinstein, P. R. & Liu, J. Collaterals: Implications in cerebral ischemic diseases and therapeutic interventions. Brain Res. 1623, 18–29 (2015).

11. Kluytmans, M. et al. Cerebral hemodynamics in relation to patterns of collateral flow. Stroke. 30, 1432–1439 (1999).

12. Seners, P. et al. Better Collaterals Are Independently Associated With Post-Thrombolysis Recanalization Before Thrombectomy. Stroke 867–872 (2019) doi: 10.1161/STROKEAHA.118.022815.

13. Shuaib, A., Butcher, K., Mohammad, A. A., Saqqur, M. & Liebeskind, D. S. Collateral blood vessels in acute ischaemic stroke: A potential therapeutic target. Lancet Neurol. 10, 909–921 (2011).

14. Harder, D. R., Alkayed, N. J., Lange, A. R., Gebremedhin, D. & Roman, R. J. Functional hyperemia in the brain: Hypothesis for astrocyte-derived vasodilator metabolites. Stroke 29, 229–234 (1998).

15. Kim, T., Masamoto, K., Fukuda, M., Vazquez, A. & Kim, S. G. Frequency-dependent neural activity, CBF, and BOLD fMRI to somatosensory stimuli in isoflurane-anesthetized rats. Neuroimage 52, 224–233 (2010).

16. Yaseen, M. A. et al. Microvascular oxygen tension and flow measurements in rodent cerebral cortex during baseline conditions and functional activation. J. Cereb. Blood Flow Metab. 31, 1051–1063 (2011).

17. Takuwa, H. et al. Hemodynamic changes during somatosensory stimulation in awake and isoflurane-anesthetized mice measured by laser-Doppler flowmetry. Brain Res. 1472, 107–112 (2012).

18. Bizeau, A. et al. Stimulus-evoked changes in cerebral vessel diameter: A study in healthy humans. J. Cereb. Blood Flow Metab. 38, 528–539 (2018).

19. Liao, L. De et al. Rescue of cortical neurovascular functions during the hyperacute phase of ischemia by peripheral sensory stimulation. Neurobiol. Dis. 75, 53–63 (2015).

20. Pan, H.-C. et al. Neurovascular function recovery after focal ischemic stroke by enhancing cerebral collateral circulation via peripheral stimulation-mediated interarterial anastomosis. Neurophotonics 4, 1 (2017).

21. Burnett, M. G. et al. Electrical forepaw stimulation during reversible forebrain ischemia decreases infarct volume. Stroke 37, 1327–1331 (2006).

22. Carmichael, S. T. Rodent models of focal stroke: size, mechanism, and purpose. NeuroRx 2, 396–409 (2005).

23. Dirnagl, U. Rodent Models Of Stroke. Neuromethods vol. 120 (2016).

24. Xu, S.-Y. & Pan, S.-Y. The failure of animal models of neuroprotection in acute ischemic stroke to translate to clinical efficacy. Med. Sci. Monit. Basic Res. 19, 37–45 (2013).

25. Clark, T. A. et al. Artery targeted photothrombosis widens the vascular penumbra, instigates peri-infarct neovascularization and models forelimb impairments. Sci. Rep. 9, 1–12 (2019).

26. Munting, L. P. et al. Influence of different isoflurane anesthesia protocols on murine cerebral hemodynamics measured with pseudo-continuous arterial spin labeling. NMR Biomed. 32, 1–12 (2019).

27. Xie, H. et al. Differential effects of anesthetics on resting state functional connectivity in the mouse. J. Cereb. Blood Flow Metab. (2019) doi: 10.1177/0271678X19847123.

28. Schlegel, F., Schroeter, A. & Rudin, M. The hemodynamic response to somatosensory stimulation in mice depends on the anesthetic used: Implications on analysis of mouse fMRI data. Neuroimage 116, 40–49 (2015).

29. Dunn, A. K. Laser speckle contrast imaging of cerebral blood flow. Ann. Biomed. Eng. 40, 367–377 (2012).

30. Kazmi, S. M. S., Balial, S. & Dunn, A. K. Optimization of camera exposure durations for multi-exposure speckle imaging of the microcirculation. Biomed. Opt. Express 5, 2157 (2014).

31. Schrandt, C. J., Kazmi, S. M. S., Jones, T. A. & Dunn, A. K. Chronic monitoring of vascular progression after ischemic stroke using multiexposure speckle imaging and two-photon fluorescence microscopy. J. Cereb. Blood Flow Metab. 35, 933–42 (2015).

32. Sullender, C. T. et al. Imaging of cortical oxygen tension and blood flow following targeted photothrombotic stroke. Neurophotonics 5, 1 (2018).

33. Armitage, G. A., Todd, K. G., Shuaib, A. & Winship, I. R. Laser speckle contrast imaging of collateral blood flow during acute ischemic stroke. J. Cereb. Blood Flow Metab. 30, 1432–1436 (2010).

34. Zhang, H., Prabhakar, P., Sealock, R. & Faber, J. E. Wide genetic variation in the native pial collateral circulation is a major determinant of variation in severity of stroke. J. Cereb. Blood Flow Metab. 30, 923–934 (2010).

35. Lay, C. C., Davis, M. F., Chen-Bee, C. H. & Frostig, R. D. Mild sensory stimulation completely protects the adult rodent cortex from ischemic stroke. PLoS One 5, (2010).

36. Frostig, R. D., Lay, C. C. & Davis, M. F. A rat’s whiskers point the way toward a novel stimulus-dependent, protective stroke therapy. Neuroscientist 19, 313–328 (2013).

37. Hancock, A. M. & Frostig, R. D. Testing the effects of sensory stimulation as a collateral-based therapeutic for ischemic stroke in C57BL/6J and CD1 mouse strains. PLoS One 12, 1–16 (2017).

38. Wu, Q. J. & Tymianski, M. Targeting nmda receptors in stroke: New hope in neuroprotection. Mol. Brain 11, 1–14 (2018).

39. Paz, J. T., Christian, C. A., Parada, I., Prince, D. A. & Huguenard, J. R. Focal Cortical Infarcts Alter Intrinsic Excitability and Synaptic Excitation in the Reticular Thalamic Nucleus. J. Neurosci. 30, 5465–5479 (2010).

40. Liao, F. et al. Murine versus human apolipoprotein E4: differential facilitation of and co-localization in cerebral amyloid angiopathy and amyloid plaques in APP transgenic mouse models. Acta Neuropathol. Commun. 3, 70 (2015).

